# An emerging PB2-627 polymorphism increases the pandemic potential of avian influenza virus by breaking through ANP32 host restriction in mammalian and avian hosts

**DOI:** 10.1101/2024.07.03.601996

**Authors:** Yuxin Guo, Sicheng Shu, Yong Zhou, Wenjing Peng, Zhimin Jiang, Yudong Li, Tian Li, Fanshu Du, Linlin Wang, Xue Chen, Jinze Dong, Chuankuo Zhao, Maggie Haitian Wang, Yipeng Sun, Honglei Sun, Lu Lu, Paul Digard, Kin-Chow Chang, Hui-Ling Yen, Jinhua. Liu, Juan Pu

## Abstract

Alterations in the PB2-627 domain could substantially increase the risk of an avian influenza virus (AIV) pandemic. So far, a well-known mammalian mutation PB2-E627K has not been maintained in AIV in poultry, which limits the spread of AIVs from avian to humans. Here, we discovered a variant, PB2-627V, which combines the properties of avian-like PB2-627E and human-like PB2-627K, overcoming host restrictions and posing a risk for human pandemics. Specifically, by screening the global PB2 sequences, we discovered a new independent cluster with PB2-627V emerged in the 2010s, which is prevalent in various avian, mammalian, and human isolates of AIVs, including H9N2, H7N9, H3N8, 2.3.4.4b H5N1, and other subtypes. And, the increasing prevalence of PB2-627V in poultry is accompanied by a rise in human infection cases with this variant. Then we systematically assessed its host adaptation, fitness, and transmissibility across three subtypes of AIVs (H9N2, H7N9, and H3N8) in different host models, including avian and human cells, chickens, mice, and ferrets where infections naturally occur. We found that PB2-627V facilitates AIVs to efficiently infect and replicate in chickens and mice by utilizing both avian- and human-origin ANP32A proteins. Importantly, and like PB2-627K, PB2-627V promotes efficient transmission between ferrets through respiratory droplets. Deep sequencing in passaged chicken samples and transmitted ferret samples indicates that PB2-627V remains stable across the two distinct hosts and has a high potential for long-term prevalence in avian species. Therefore, the mutation has the ability to continue spreading among poultry and can also overcome the barrier between birds and humans, greatly enhancing the likelihood of AIVs infecting humans. Given the escalating global spread of AIVs, it is crucial to closely monitor influenza viruses carrying PB2-627V to prevent a pandemic.

## Introduction

Avian influenza viruses (AIVs) are critically important in the evolution of pandemic influenza viruses [1]. Adaptive changes in AIVs allow breakthrough infection in mammalian species through altering viral interactions with host-specific factors [2–4]. The current clade 2.3.4.4b highly pathogenic avian influenza (HPAI) H5Ny virus outbreaks have swept across continents, posing an unprecedented panzootic threat to birds, mammals, and public health [5] (https://www.who.int/news/item/12-07-2023-ongoing-avian-influenza-outbreaks-in-animals-pose-risk-to-humans). Other subtypes of AIVs have also been responsible for recurrent human infections [6]. For example, H7N7 [7], H7N9 [8,9], H10N8 [10], H5N6 [11], H10N3 [12] and H3N8 [13] viruses have caused over 1600 documented human cases [11]. Notably, most of these AIV subtypes possess internal (i.e. non-glycoprotein) genes that are derived from avian H9N2 viruses [8,11–13]. H9N2 virus is globally widespread in wild and domestic birds [14,15], and causes repeated human infections, especially since 2015. In these interspecies transmission events, critical adaptive mutations often occur on the PB2 protein of AIVs [12,16–19].

PB2 is one of three polymerase subunits of the influenza virus RNA-dependent RNA polymerase (RdRP), crucial for genome transcription and replication [20]. Specific amino acid changes have been identified in PB2 as key determinants of avian-to-mammalian adaptation, through its interaction with host acidic leucine-rich nuclear phosphoprotein 32 (ANP32) in avian and mammalian cells [21–24]. Their interactions exhibit host-species specificity [23]. PB2-627K in human influenza viruses interacts with human ANP32A and ANP32B (huANP32A and huANP32B) to support viral polymerase function [22]. On the other hand, PB2-627E in AIV can utilize avian ANP32A, but not huANP32A or huANP32B [22]. PB2-E627K amino acid substitution in AIVs promotes viral polymerase activity in mammalian cells by enhancing PB2 interaction with huANP32A and huANP32B [25]. Fortunately, the PB2-627K genotype is only frequently detected in AIVs isolated from human infections, and rarely detected in AIVs isolated from poultry or wild birds [26], greatly reducing the possibility of AIVs transmission to humans. In this study, we discovered an emerging PB2-627 polymorphism (PB2-627V) in diversified subtypes that increases the pandemic potential of AIVs by breaking through ANP32 host restriction in mammalian and avian hosts.

## Results

### PB2-627V prevalence in influenza virus isolates from avian and mammalian species

Given the crucial role of PB2 residue 627 in host adaptation, we analyzed 42,297 influenza PB2 sequences worldwide (excluding human H3N2 and H1N1pdm09 viruses) for polymorphisms at this position. We found that the number of sequences with PB2-627V rapidly increased after 2016 (Fig 1A). Among these viruses with PB2-627V, 71.3% were isolated from avian species, 14.7% were from humans, 14.0% were from other mammalian hosts (such as fox, swine, tiger, and mink) and environment (Fig 1A and S1 Table). In geographical distribution, 61.7% of the viruses were isolated in China, with the remaining (38.3%) found in the Middle East, Germany, Mexico, Canada, Italy, Spain, Russia, America, Denmark, and Belgium (Fig. 1A, and S1 Table). In terms of subtype distribution, 64.5% of PB2-627V variants were detected in H9N2 viruses, 9.9% in H7N9 and 9.1% in H5N6 viruses; the remaining 16.5% were in various other subtypes: H1N1, H1N2, H3N2, H3N8, H5N1, H5N2, H6N6, H7N1, and H10N3 (Fig. 1A, and S1 Table). Notably, PB2-627V has also been identified in clade 2.3.4.4b H5N1 viruses isolated from wild birds and red foxes (Fig. 1A, and S1 Table).

**Fig 1.**
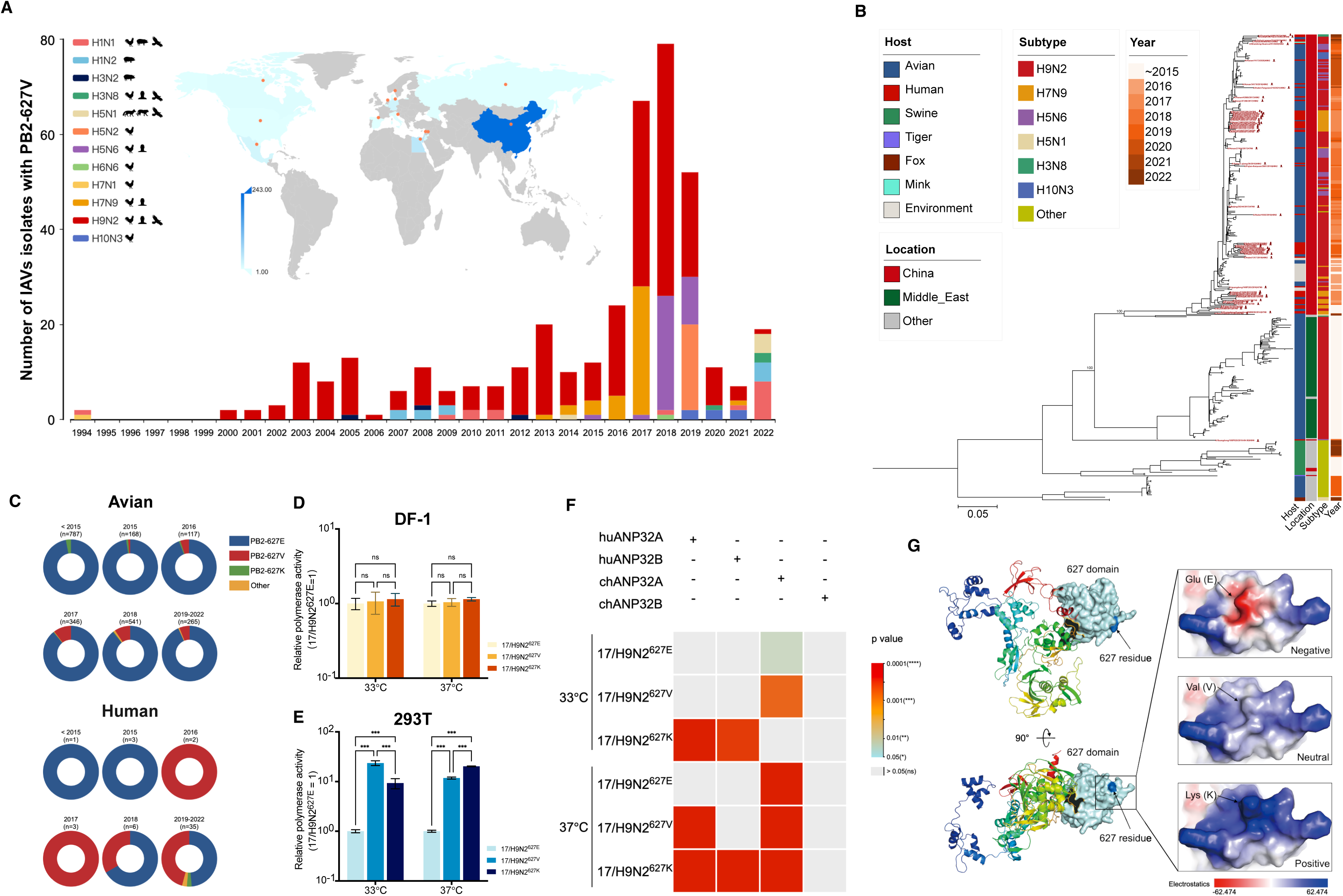
Emergence and cross-host species function of PB2-627V. (**A).** The global prevalence of PB2-627V in influenza A viruses (IAVs) (excluding human H3N2 and H1N1pdm09 viruses) during 1994-2022. Bars show the number of isolates by subtype and host, while a world map displays the isolate’s number by location. (**B).** Phylogenetic tree of PB2 genes with the residue 627V collected worldwide during 1994-2022 (n=394). Human isolates’ names are in red. (**C).** Amino acid polymorphisms of the PB2-627 residue in avian and human-derived H9N2 viruses isolated in China. All sequence data were collected from the GISAID database. (**D-E)**. Relative polymerase activities at 33°C or 37°C. DF-1 cells **(D)** and 293T cells **(E)** were transfected with firefly minigenome reporter, renilla expression control, and polymerase (PB2, PB1, PA, and NP) of 17/H9N2^627E^, 17/H9N2^627V^, or 17/H9N2^627K^. Cells were incubated for 24 hours and the luciferase activities, normalized to corresponding renilla luciferase activity, are presented relative to 17/H9N2^627E^ (set as 1). Each data point is the mean ± SD of three independent experiments (one-way ANOVA). **(F).** Heat map shows p-values of polymerase activities in DKO cells overexpressing ANP32 proteins. chANP32A, chANP32B, huANP32A, huANP32B, or empty vector was transfected into DKO cells, incubated at 33°C and 37°C for 24 hours, and assayed the luciferase activity. Statistical significance was based on one-way ANOVA and subsequent Dunnett’s test, compared with the corresponding value of empty vector (ns, not significant; *, p<0.05; **, p<0.01; ***, p<0.001). (**G).** Impact of amino acids at the PB2-627 position on the surface charge of the PB2 protein. Red (negative) and blue (positive) proteins are shown.

In phylogenetic analysis, recent PB2-627V sequences mainly belong to a major group that includes most human and avian isolates from multiple subtypes (H9N2, H7N9, H5N6, H3N8, H10N3, and H5N1 viruses) (Fig 1B). For H9N2 subtype, PB2 gene showed the evolution of a PB2-627V cluster (S1A Fig). Ancestral host reconstruction on time-scaled phylogeny indicated that chicken isolates are the origin of human isolates within this cluster (S1B Fig). Notably, PB2-627V of H9N2 is more frequently detected in humans than avian isolates from 2016-2022 (Fig 1C). Similarly, there was a dramatic increase in PB2-627V detection in avian and human hosts for the H7N9, H5N6, and H3N8 subtypes after 2016 (S2 Fig). Thus, it appears that PB2-627V has achieved better adaptability in poultry populations and its emerging prevalence both in avian and humans suggests functional significance across various host species.

### PB2-627V promotes polymerase activity of H9N2 AIV in human and chicken cells by utilizing ANP32 proteins of both host species

To evaluate the functionality of PB2-627V, we used a previously identified chicken H9N2 isolate A/chicken/Hubei-m0530/2017 (referred to as 17/H9N2^627E^) with PB2-627E as the backbone, and introduced the PB2-E627V and PB2-E627K changes. Although PB2-627V and - 627K are found in other natural chicken isolates, this approach allowed the dissection of the function of this single mutation. A dual-luciferase assay system was used to compare viral polymerase activities conferred by PB2-627E, V, and K in avian DF-1 cells and human 293T cells at 33°C and 37°C, respectively, mimicking the temperature of human upper and lower respiratory tract [27]. In DF-1 cells, 17/H9N2^627E^, 17/H9N2^627V^, and 17/H9N2^627K^ showed comparable polymerase levels at 33°C and 37°C (Fig 1D). However, a 23.6-fold (±2.6) increase in polymerase activity at 33°C, and an 11.8-fold (±0.6) increase at 37°C were found with 17/H9N2^627V^ in 293T cells compared with 17/H9N2^627E^ (p < 0.001); 17/H9N2^627K^ exhibited a 9.3-fold (±2.6) increase in polymerase activity at 33°C and a 20.2-fold (±0.4) increase at 37°C (p < 0.001) relative to 17/H9N2^627E^ (Fig. 1E). Accordingly, in avian DF1 cells, PB2-627V demonstrated more robust viral transcriptional and replication functions than PB2-627K, while in human A549 cells, it surpassed PB2-627E (S3A and S3B Fig).

Viral polymerase activity was further examined in ANP32A and ANP32B double-knockout 293T (DKO) cells over-expressing chicken- or human-ANP32 protein(s) [25]. As expected, polymerase activities of all 3 viruses (17/H9N2^627E^, 17/H9N2^627V^, and 17/H9N2^627K^) in control DKO cells were significantly lower than those in the wild type 293T cells (S3C Fig). Polymerase activity of 17/H9N2^627E^ in the DKO cells was significantly enhanced in the presence of chicken ANP32A (chANP32A) at both 33°C and 37°C (p < 0.05) (Fig 1F). With 17/H9N2^627V^, chANP32A in DKO cells significantly enhanced the polymerase activity at both 33°C and 37°C (p < 0.05), and huANP32A significantly enhanced polymerase activity only at 37°C (p < 0.001) (Fig 1F). With 17/H9N2^627K^, expression of chANP32A in DKO cells significantly enhanced viral polymerase activity only at 37°C (p < 0.001), and huANP32A and huANP32B significantly enhanced the polymerase activity at both 33°C and 37°C (p < 0.001) (Fig 1F).

ANP32 interface with PB2 at the 627-domain by electrostatic and hydrophobic interactions [28]. PB2-627K (lysine) has a positive charge and PB2-627E (glutamic acid) has a negative charge. Modeling the surface electrostatic potential predicted that valine at PB2-627 confers a neutral state (Fig 1G), which could permit the PB2 protein to interact sufficiently with avian and human ANP32A proteins to promote viral polymerase activity in avian and human cells. In summary, these findings indicate that PB2-627V can utilize chicken and human ANP32A in the respective species of cells to facilitate viral polymerase activity.

### PB2-627V in H9N2 virus maintains efficient replication in chickens and facilitates replication in human cells and mice

Next, viruses with the three PB2 627 polymorphisms were rescued on the background of the 17/H9N2 virus. The replication kinetics of these 17/H9N2^627E^, 17/H9N2^627V^, and 17/H9N2^627K^ viruses were assessed in avian CEF cells; no significant differences in replication were found between viruses (p > 0.05) (Fig 2A). Chickens individually inoculated with 10^6^ EID_50_ of each virus showed no mortality (S4A Fig) and no apparent clinical signs. Viral loads detected from oropharyngeal swabs of 17/H9N2^627V^ and 17/H9N2^627E^ inoculated chickens were significantly higher than those infected with 17/H9N2^627K^ at 6 and/or 8 dpi (p < 0.05) (Fig 2B). The three viruses replicated to comparable titers in the turbinates at 3 dpi, but 17/H9N2^627V^ replicated to higher titers than 17/H9N2^627K^ in the lungs (p < 0.05) (Fig 2C). Histopathological examination of chicken lungs at 3 dpi indicate that all viruses induced inflammatory responses, with more hemorrhage noted in 17/H9N2^627V^ infected lungs (Fig 2D). These findings indicate that PB2-627V and PB2-627E replicated comparably well in chickens.

**Fig 2.**
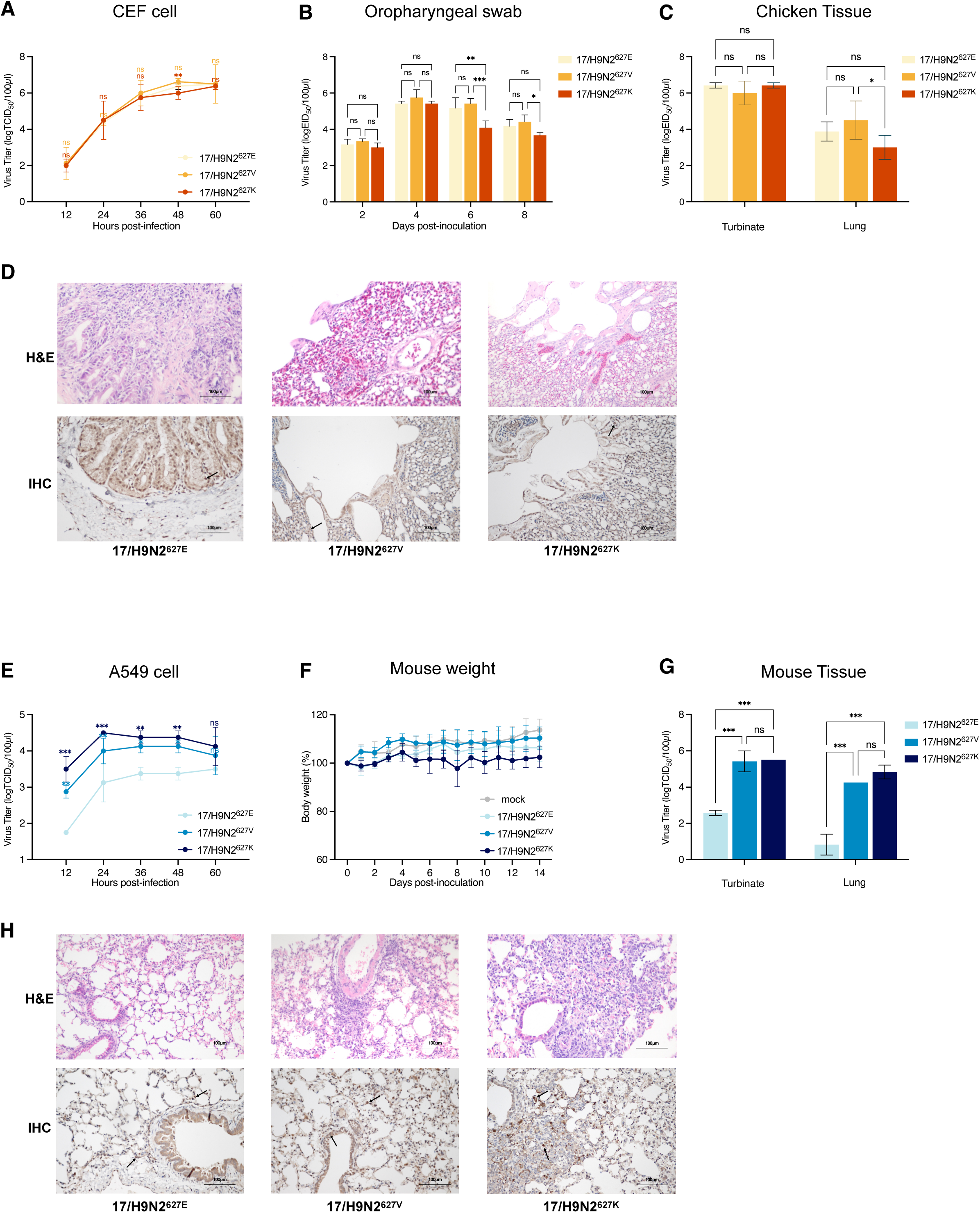
PB2-627V conferred dual adaptability to H9N2 virus to replicate in avian and mammalian species. (**A).** CEF cells were infected with indicated viruses at 0.01 MOI and virus titers were determined at indicated times post-infection by TCID_50_ assays. (**B-D).** Groups of 6 chickens were i.n. infected with indicated viruses at 10^6^ EID_50_, three chickens from each group were used to determine the viral titers (**B**) from oropharyngeal samplings at 2, 4, 6, and 8 dpi. Another three chickens from each group were euthanized at 3 dpi. Virus titers (**C**) of the turbinates and lungs were determined. Representative histopathological findings (**D**) in the lungs of infected mice. Lung sections were stained with H&E (upper) and by IHC against influenza viral NP antigen (lower). Scale bars, 100 mm. (**E).** A549 cells were infected with indicated viruses at 0.1 MOI and virus titers were determined by TCID_50_ assays at indicated times post-infection. (**F-H).** Groups of 8 mice were infected with the indicated virus at doses of 10^6^ TCID_50_. Five mice of each group were monitored for body weight changes (**F**) over 14 days. Additional three mice from each group at 3 dpi were euthanized. Virus titers (**G**) in the turbinates and lungs were determined by TCID_50_ assays. Representative histopathological findings (**H**) in the lungs of infected mice. Lung sections were stained with H&E (upper) and by IHC against influenza viral NP antigen (lower). Scale bars, 100 mm. Data are represented as mean ± SD. Statistical significance was based on two-way ANOVA, compared with the corresponding value of 17/H9N2^627E^ (ns, not significant; *, p < 0.05; **, p < 0.01; ***, p < 0.001).

In infected human A549 cells, virus titers of 17/H9N2^627V^ and 17/H9N2^627K^ were significantly higher than that of 17/H9N2^627E^ from 12 to 48 hpi (p < 0.001); there was no significant difference between 17/H9N2^627V^ and 17/H9N2^627K^ (p > 0.05) (Fig 2E). Mice inoculated with 17/H9N2^627E^, 17/H9N2^627V^, or 17/H9N2^627K^ virus, showed no obvious clinical signs, as in infected chickens (Fig 2F and S4B Fig). Higher virus loads were detected in the nasal turbinates and lungs of mice infected with 17/H9N2^627V^ and 17/H9N2^627K^ at 3 dpi than those with 17/H9N2^627E^ (p < 0.001); there was no significant difference in virus titers between the 17/H9N2^627V^ and 17/H9N2^627K^ groups (p > 0.05) (Fig 2G). Likewise, 17/H9N2^627V^ and 17/H9N2^627K^ viruses produced more viral NP protein-positive cells in the lung than in corresponding 17/H9N2^627E^ infected mice (Fig 2H). Thus, the infectivity of PB2-627V H9N2 virus in human cells and mice is similar to that of PB2-627K virus.

### PB2-627V confers aerosol transmission of H9N2 virus in ferrets

Transmissibility of influenza viruses between ferrets is an indicator of potential epidemic or pandemic risk. Direct contact (DC) and respiratory droplet (RD) transmission tests of the 17/H9N2^627E^, 17/H9N2^627V^, and 17/H9N2^627K^ viruses were performed in ferrets. All viruses infected the directly infected donor animals, as assessed by virus titers in nasal washes (Fig. 3A-3C, and S2 Table). 17/H9N2^627V^ and 17/H9N2^627K^ viruses were transmitted to all DC ferrets and all RD ferrets, but 17/H9N2^627E^ was only transmitted to DC groups. Transmission by RD in the 17/H9N2^627V^ group took place around 2 days later than in the 17/H9N2^627K^ group (Fig 3A-3C). Areas under the curve (AUC), based on virus titers over time in the nasal washes, were determined to compare total viral shedding during infection (Fig 3D and S3 Table). In donor ferrets, virus production was comparable between the three viruses. In DC ferrets, 17/H9N2^627V^ virus output (AUC = 18.13 ± 3.69) was slightly higher than that of 17/H9N2^627E^ virus (14.13 ± 3.01, p = 0.19) and 17/H9N2^627K^ virus (16.25 ± 1.27, p = 0.68). In RD ferrets, 17/H9N2^627V^ virus output (AUC = 11.75 ± 2.88) was slightly lower than that of 17/H9N2^627K^ virus (15.42 ± 1.60, p = 0.25) (Fig 3D and S3 Table). Thus, in contrast to PB2-627E, PB2-627V promotes aerosol transmissibility of H9N2 virus in ferrets.

**Fig 3.**
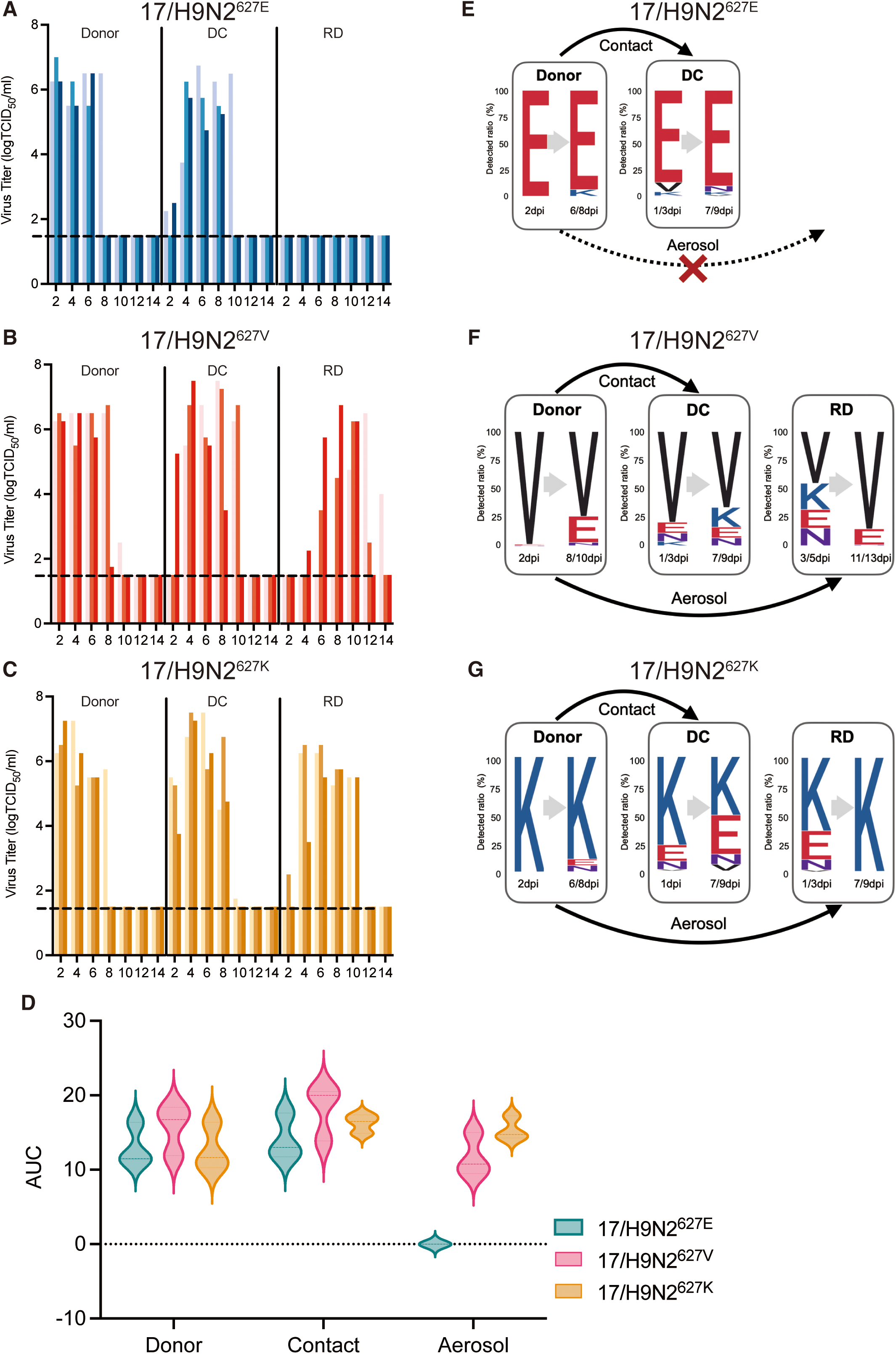
PB2-627V facilitated aerosol transmissibility of H9N2 virus in ferrets. Each ferret (n = 3) was inoculated with 10^6^ TCID_50_ of indicated virus. The next day, the infected ferret was individually paired by co-housing with a DC ferret; an RD-contact ferret was also housed in a wire frame cage adjacent to the infected ferret. Nasal washes were collected for virus-shedding detection every other day from all ferrets from 2 dpi. (**A-C).** Virus titers of nasal washes were determined by TCID_50_ assays on MDCK cells. Each color bar represents virus titers of an individual ferret. Dashed lines indicate the lower limit of virus detection (10^1.5^ TCID_50_/ml). (**D).** AUC values approximate total virus output throughout infection (mean AUC from three ferrets). (**E-G).** For each infected or transmitted ferret, the nasal washes collected at the first and last virus detection time were analyzed by deep sequencing. Detection time points are based on the results shown in Fig. 3, A-C. The height of each letter indicates the relative ratio of amino acids at PB2-627. Sequencing was performed on an Illumina NovaSeq platform.

To monitor any polymorphic changes at residue PB2-627 during virus infection and transmission, next-generation sequencing was performed on nasal wash samples from each ferret at the first and the last time points of positive virus detection. In 17/H9N2^627E^ groups, 627E remained the dominant genotype in the donors and DC ferrets (Fig 3E). In the donor groups of 17/H9N2^627V^ and 17/H9N2^627K^, 627V and 627K were the corresponding dominant genotypes. In the DC and RD transmission groups of 17/H9N2^627V^ and 17/H9N2^627K^, polymorphisms at PB2-627 during infection were more noticeable, but at the last detection time point of each RD group, corresponding 627V and 627K returned to their earlier dominance (Fig 3F and 3G). These results suggest that residue PB2-627V stably confers aerosol transmission of H9N2 virus in ferrets.

### PB2-627V has the potential for sustained prevalence in poultry

17/H9N2^627E^, 17/H9N2^627V^, and 17/H9N2^627K^ H9N2 viruses were serially passaged in chickens to assess the potential for sustained prevalence of PB2-627V in poultry. Chickens were directly infected with each virus (Passage 1, P1) and subsequently (P2 - P5) passaged through contact transmission (Fig 4A). All viruses were successfully passaged from P1 to P5. We then performed next-generation sequencing of whole viral genomes from RNA isolated from the lungs of the P5 chickens. Mutations above 1% threshold were identified as intra-host single nucleotide variants (iSNVs) [29,30]. Overall, most iSNVs were detected at low frequency and there were no significant differences between the 3 groups (Fig 4B). Notably, variant frequency at residue PB2-627 was below the 1% threshold in all P5 chicken samples (Fig 4C). At the amino-acid level, two variant residues (PA-D529N, NP-T211D) occurred at a high frequency (>10%) in the 17/H9N2^627K^ group, but no high-frequency variant residues were detected in 17/H9N2^627E^ and 17/H9N2^627V^ groups (Fig 4D), suggesting that compared to PB2-627E and PB2-627V, PB2-627K required other cooperative mutations in RNP genes for transmission in chickens. Taken together, PB2-627V showed a high potential for sustained prevalence in chickens.

**Fig 4.**
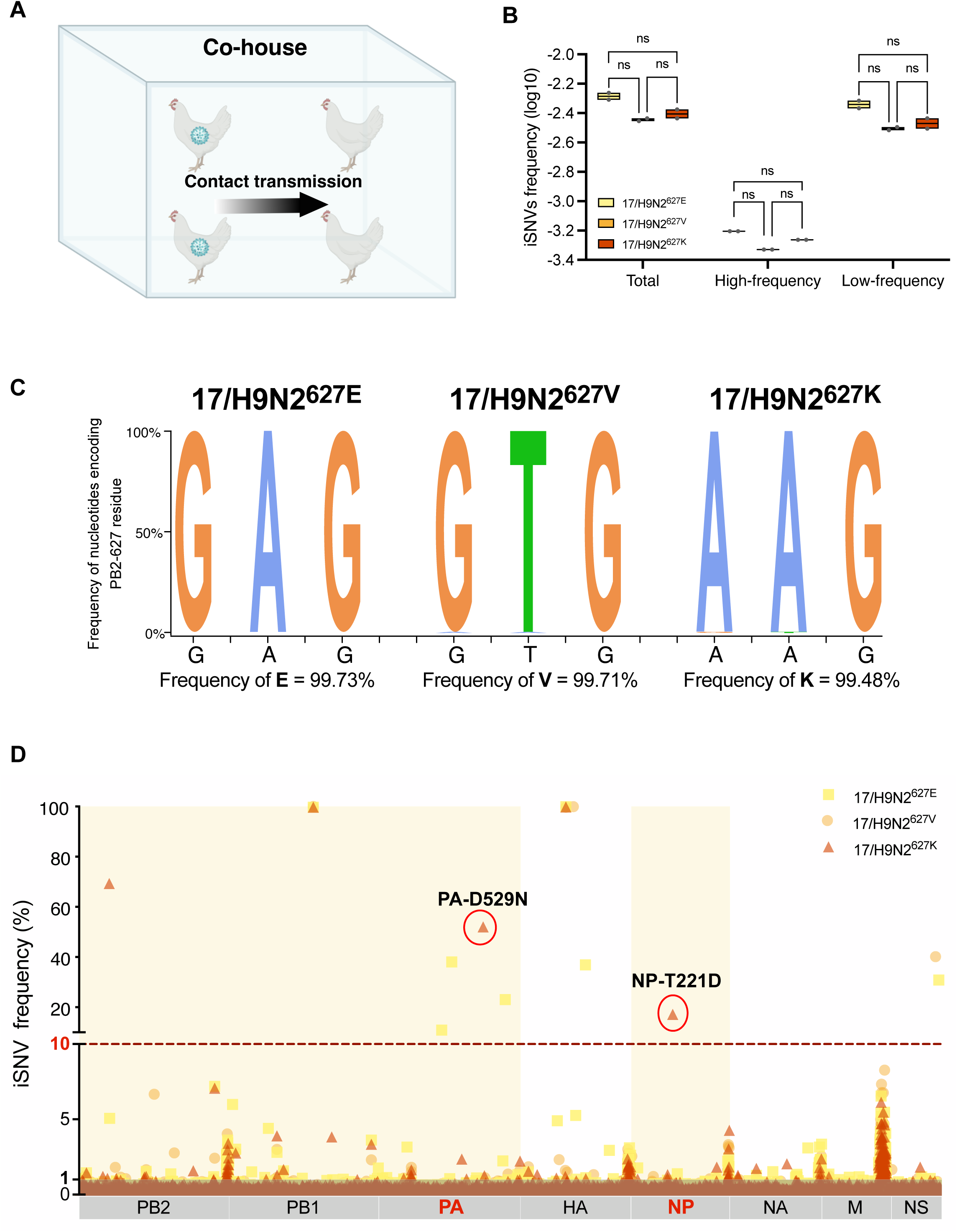
PB2-627V has the potential for sustained prevalence in poultry. **(A).** Schematic depiction of virus passage in chickens. Groups of 2 chickens were each i.n. infected with 10^6^ TCID_50_ of indicated virus (P1), and subsequently passaged (P1-P5) by contact. (**B).** Intra-host single nucleotide variants (iSNVs) frequency of P5 chicken. Variants of nucleotide site detected above 1% threshold were identified as iSNVs. All iSNVs were further determined as high-frequency (>10%) and low-frequency (≤10%). iSNVs frequency of the whole genome was calculated by using the number of iSNVs divided by the total number of genome sites. iSNVs frequency was expressed as the log. (**C)**. Variant frequency nucleotides encoding PB2-627 residue. The height of each letter indicates the average ratio of nucleotides encoding PB2-627 residue in two samples of each group. (**D)**. Graphs represent all iSNVs detected in P5 chicken. Variant frequencies are plotted by genome location and are colored by virus groups. two high-frequency non-synonymous iSNVs are circled and labeled with the corresponding amino acid mutations. Yellow backgrounds depict RNP genes (PB2, PB1, PA, and NP). Sequencing was performed on an Illumina NovaSeq platform. Data are presented as mean ± SD. Statistical significance was based on two-way ANOVA (ns, not significant, *, p < 0.05; **, p < 0.01; ***, p < 0.001).

### PB2-627V confers dual ability to replicate in chickens and mice to avian H7N9 and H3N8 viruses

We determined if the dual host adaptability of PB2-627V, detected at high frequency in the H9N2 virus, is transferrable to other AIV subtypes. H7N9 and H3N8 viruses possess the PB2 gene and other internal genes derived from the H9N2 viruses [9,31]. The natural mutations PB2-E627V and PB2-E627K were separately introduced into the LPAI H7N9 (A/chicken/China/0606-12/2017, referred as 17/H7N9^627E^) and H3N8 (A/chicken/Anhui/FE12/2022, referred as 22/H3N8^627E^) [32] viruses. The resulting PB2-627V, PB2-627K, and PB2-627E viruses were assessed for replication and pathogenicity in chickens and mice.

In chickens, introduction of PB2-E627V or PB2-E627K mutation into 17/H7N9 and 22/H3N8 viruses did not significantly affect replication or pathogenicity relative to their PB2-627E counterparts (Fig 5A-5E), although the replication of 17/H7N9^627V^ in chicken lungs was significantly higher than that of 17/H7N9^627K^ (p<0.05) (Fig 5B). Thus, PB2-E627V had little effect on virus replication and pathogenicity in chickens (Fig. 5A-5E, and S5A Fig). In mice, both PB2-E627V and PB2-E627K changes in 17/H7N9 and 22/H3N8 viruses led to virus loads in the nasal turbinates and lungs, weight loss, and mortality (p<0.01) (Fig 5F-5K). Murine lungs at 3 dpi. with subtypes housing PB2-627V showed more severe pathology than those with PB2-627E (Fig 5L for 17/H7N9 and S5B Fig for 22/H3N8). In summary, PB2-E627V significantly increases pathology and replication of H7N9 and H3N8 virus subtypes in mice and maintains their replication fitness in chickens.

**Fig 5.**
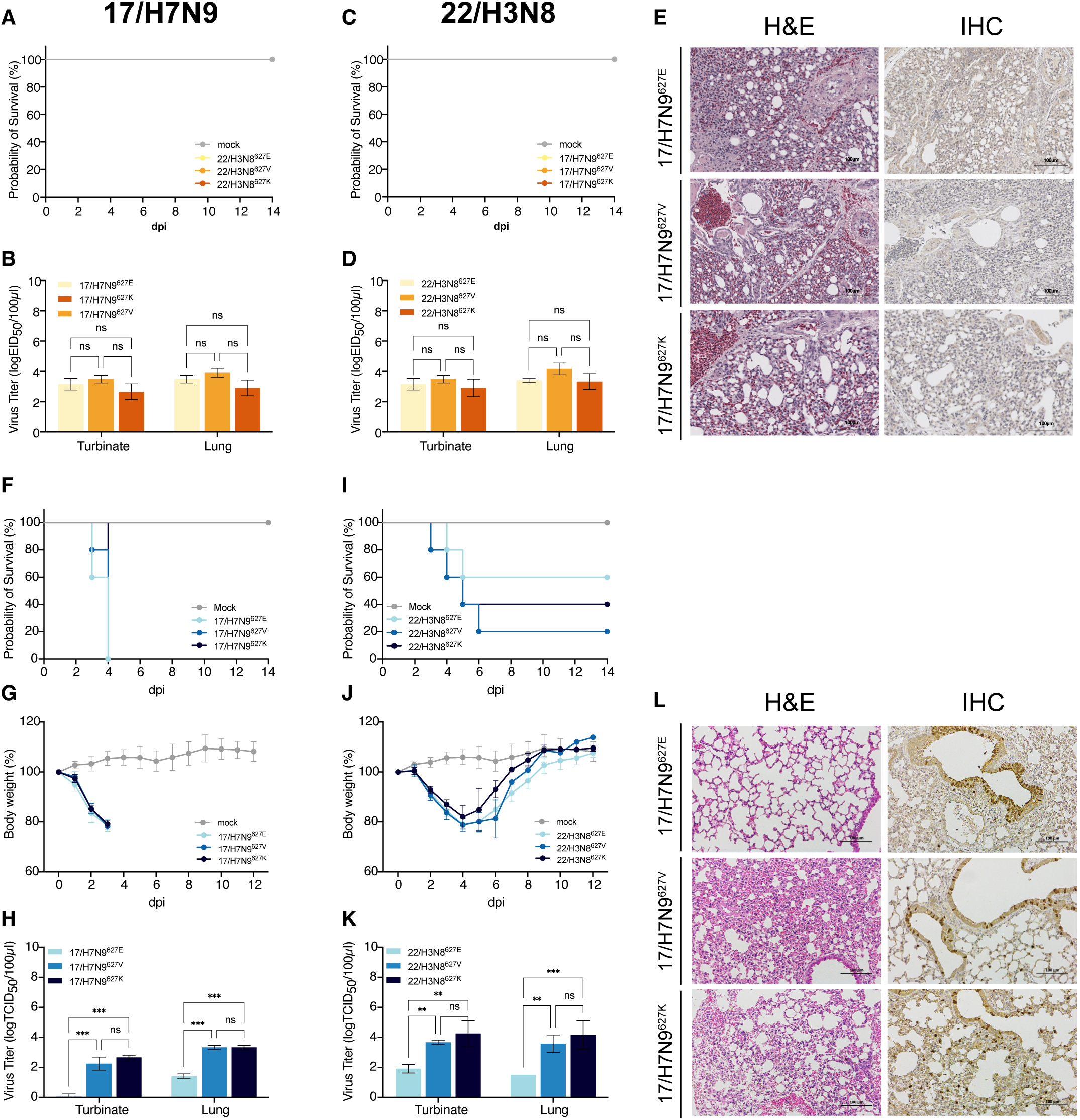
PB2-627V conferred dual adaptability in avian H7N9 and H3N8 viruses to replicate in chickens and mice. **(A-E).** Groups of 6 chickens were infected with indicated viruses at doses of 10^6^ EID_50_. Three infected chickens in each group were monitored for survival (**A**, **C)** over 14 days. At 3 dpi, the other three chickens in each group were euthanized. Virus titers (**B**, **D**) in the turbinates and lungs were determined by TCID_50_ assays. Representative histopathological findings (**E**) in the lungs of infected mice. Lung sections were stained with H&E (left) and by IHC against influenza viral NP antigen (right). Scale bars, 100 mm. (**F-L).** Groups of 8 mice were infected with indicated viruses at doses of 10^6^ TCID_50_. Five mice in each group were monitored for survival (**F**, **I**) and body weight changes (**G**, **J**) over 14 days. Mortality from infection included mice euthanized at a weight loss of 25% or more. At 3 dpi, the other three mice in each group were euthanized. Virus titers (**H**, **K**) in the turbinates and lungs were determined by TCID_50_ assays. Representative histopathological findings (**L**) in the lungs of infected mice. Lung sections were stained with H&E (left) and by IHC against influenza viral NP antigen (right). Scale bars, 100 mm. Data presented are represented as mean ± SD. Statistical significance was based on two-way ANOVA (ns, not significant; *, p < 0.05; **, p < 0.01; ***, p < 0.001).

## Discussion

Here, we identified PB2-627V as an emerging variant in AIVs with significant zoonotic and pandemic potential. We observed that PB2-627V enhances the polymerase activity of H9N2 AIV in both human and chicken cells by utilizing ANP32A proteins from both species, thereby overcoming the restriction imposed by human ANP32 on AIVs carrying PB2-627E and exhibiting properties similar to those with PB2-627K. PB2-627V in H9N2 virus maintains efficient replication in chickens, facilitates virus replication in mice, and confers aerosol transmission in ferrets. Furthermore, PB2-627V is stably maintained in passages through chickens and also conferred the dual ability to replicate in chickens and mice to avian H7N9 and H3N8 subtype viruses. Thus, the increased prevalence of this dual adaptive mutation of PB2-E627V in poultry greatly increases the chances of AIVs infecting humans. In addition, the wide distribution of this mutation in multiple subtypes, hosts, and countries poses global public health risks.

Clinical data and phylogenetic analysis revealed that residue PB2-627V originated in chicken influenza viruses (likely the H9N2 subtype) and was subsequently transmitted to humans. The prevalence of PB2-627V in AIVs is on the rise, particularly within the H9N2 subtype, and PB2-627V is also found in other subtypes such as H7N9, H5N6, H3N8, and H10N3, which contain reassorted internal genes derived from H9N2 [8,11–13]. Serial passage studies in chickens further indicate that additional mutations occurred in the polymerase protein after the passage of H9N2 viruses with the PB2-E627K. However, H9N2 viruses with PB2-627E and -627V did not produce further mutations. This may be due to the PB2-627K mutation still having defects in transmission in chickens and requiring further adaptation. This could explain why PB2-627K is very rare in avian isolates. The PB2-E627K mutation usually takes place in infected humans [33]. Conversely, PB2-627V appears to be a product of evolution in H9N2 virus and has a high potential for long-term prevalence in poultry populations, posing an increasing likelihood of viral transmission to humans.

Cellular ANP32 proteins specifically support influenza virus genome replication. ANP32A can bridge PB2 dimers and form extensive contacts with two polymerases to maintain a dimeric assembly [22,34]. The C-terminal low complexity acidic region (ANP32A^LCAR^) inserts between two PB2-627 domains of the polymerase dimer. Human ANP32A^LCAR^ contains only acidic amino acid residues. Thus, with human ANP32A, basic or uncharged residues (such as lysine, K or valine, V) are preferred in the PB2-627 domain to acidic amino acids (such as glutamic acid, E) [21,22]. In avian cells, the additional 33 amino acids in chANP32A^LCAR^ allow AIVs carrying PB2-627E to effectively utilize chANP32A in avian cells [23,35]. Indeed, in the context of H9N2, we found that the polymerase activity of PB2-627V in avian cells is comparable to PB2-627E, and in human cells is comparable to PB2-627K which suggests that the nonpolar neutral amino acid of PB2-627V has the dual ability to interact with avian ANP32A and human ANP32A in different host cells. It is noteworthy that, when the PB2-627 site is changed from E to V, the virus can not only undergo contact transmission in ferrets but also undertake effective aerosol transmission, similar to that of the 17/H9N2^627K^ variant. In addition, PB2-627V can be maintained in ferret post-infection, during contact and aerosol transmission. Thus, these features suggest that the dual adaptive PB2-627V AIVs reduce the host barrier between avians and humans.

The PB2-E627V has been identified in H5N1 viruses where it was shown to increase viral replication in mammalian cells and virulence in mice [36], while in the background of a laboratory strain of H1N1 (PR8), it reduced viral replication in mammalian cells compared with PB2-627K [37]. However, because the PB2-627V variant was rarely found in natural influenza isolates at that time (before 2015), it did not attract much attention. In H9N2 AIVs, PB2-627V variants appeared in the Middle East from 2000 to 2015 but were not stably maintained. The genotype of Middle East H9N2 viruses was very different from the G57 lineage [8] H9N2 virus prevalent in China and studied here [38,39]. Under experimental conditions, PB2-627V in the Middle East H9N2 virus needs to operate in concert with PB2-E543D, PB2-A655V and PB2-K526R to enhance viral replication in mammals [38]. Numerous adaptive mutations have recently been identified in the G57 H9N2 virus, including PB2-292, PB2-588, PA-356, M1-37 and HA-226 [15,40,41], in avian and human isolates [42–45]. It is plausible that one or more of these adaptive changes in the G57 genotype exert epistatic effects to induce the emergence of the stable PB2-627V residue, but this remains to be tested.

Collectively, our results suggest that the recent evolved variant PB2-E627V combines the characteristics of avian-like PB2-627E and human-like PB2-627K. It exhibits a crucial dual-adaptation that can effectively infect and transmit in both avian and mammalian hosts, thereby overcoming the interspecific barrier between birds and humans. The presence of PB2-627V in multiple subtypes of AIVs undoubtedly escalates the risk of an avian influenza virus pandemic. We therefore recommend considering PB2-627V as a new molecular marker to assess the pandemic potential of AIVs.

## Materials and Methods

### Ethics statement

9 to 11-day-old specific pathogen-free (SPF) embryonated chicken eggs and all experimental animals were purchased from Boehringer Ingelheim, China. All animal research was approved by the Beijing Association for Science and Technology (approval ID SYXK, Beijing, 2020-0053) and performed in compliance with the Beijing Laboratory Animal Welfare and Ethics guidelines, as issued by the Beijing Administration Committee of Laboratory Animals, and by the China Agricultural University (CAU) Institutional Animal Care and Use Committee guidelines (ID: SKLAB-B-2010-003) approved by the Animal Welfare Committee of CAU. The H9N2, H7N9, and H3N8 wild-type viruses used are all low pathogenic avian influenza (LPAI) viruses isolated from chickens. PB2-E627V and PB2-E627K are natural mutations found in these viruses.

### Sequence collection and alignment

The sequences of the PB2 gene were obtained from the GISAID database (http://www.gisaid.org). After removing the PB2 sequences of human H3N2 and H1N1pdm09 viruses and duplicate sequences, a total of 42,297 full-length PB2 sequences of IAVs published globally from 1994 to 2022 were obtained. All accession codes of public sequences can be found in Dataset S1. Sequences were aligned using MUSCLE v3.7 [46] via the CIPRES Science Gateway [47].

### Phylogenetic analyses

The phylogenetic tree of the H9N2 PB2 gene was constructed using a total of 4523 PB2 sequences from H9N2 subtype IAVs. The phylogenetic tree of PB2-627V was constructed using a total of 394 PB2 sequences carrying the PB2-627V. IQ-TREE version 1.6 was used to construct the maximum likelihood phylogenetic trees, applying the best-fit general time-reversible model of nucleotide substitution with gamma-distributed rate variation among sites (GTR + I + G), and performing ultrafast bootstrap resampling analysis (1000 replications). Phylogenetic trees were visualized and annotated using FigTree version 1.4.4, Adobe Illustrator 2021, and Interactive Tree Of Life (iTOL) [48].

### Host transmission dynamics analyses

A maximum clade credibility (MCC) tree was performed on the PB2-627V cluster sequences (Fig. S1A) with BEAST v1.10.4. Using the coalescent Bayesian Skygrid model, the General Time Reversible (GTR) model of evolution and the strict clock were employed. The host jump patterns were interfered by mapping the host types onto the time-scaled tree using an asymmetric discrete trait model [49]. A Markov Monte Carlo (MCMC) chain is run for 30 million, sampled every 3000 and the first 10% of tree was removed as burn-in. Stationarity and mixing were checked with Tracer v1.7.2. The MCC tree was finally generated by TreeAnnotator and visualized using FigTree v1.4.4.

### PB2 protein structure simulation

A simulation of the A/chicken/Hebei/m0530-1/2017 (17/H9N2) PB2 protein structure was performed by using Swiss-Model (https://swissmodel.expasy.org). The 17/H9N2 PB2 protein sequence was inputted in FASTA format, and the published H7N9 protein structure (PDB: 7qtl.1.C) with 98.94% sequence identity was utilized as the template. The quality of the resulting model was evaluated using GMQE and QMEAN scores. PyMOL was utilized to introduce the PB2-E627V and PB2-E627K mutations into the model and to ascertain the surface charge of the protein.

### Cell lines

Human lung epithelial cell (A549) cells, kidney epithelial (293T) cells, chicken fibroblast (DF-1) cells, and Madin-Darby Canine Kidney (MDCK) cells were maintained in our laboratory. Chick embryo fibroblast (CEF) cells were obtained from SPF embryonated chicken eggs as previously described [50]. 293T cell line with a double knockout of ANP32A and ANP32B (DKO cell) was kindly provided by Xiaojun Wang from Harbin Veterinary Research Institute. Cells were cultured with Dulbecco’s modified Eagle medium (DMEM; Gibco, New York, USA) supplemented with 10% fetal bovine serum (FBS; Gibco, Sydney, Australia) and 1% penicillin-streptomycin solution (MACGENE, Beijing, China).

### Influenza viruses

H9N2 AIV A/chicken/Hebei/m0530-1/2017 (17/H9N2) was characterized and maintained in our laboratory. Influenza H7N9 virus A/chicken/China/0606-12/2017 (17/H7N9) and influenza H3N8 virus A/chicken/Shandong/FE12/2022(H3N8) (22/H3N8) were isolated by our laboratory. Viruses were propagated in 9 to 11-day-old SPF embryonated chicken eggs at 35°C for 72 hours. Virus titers were determined by 50% tissue culture infectious dose (TCID_50_) on MDCK cells or 50% egg infectious dose (EID_50_) on SPF chicken eggs.

### Animals

5 to 6 weeks-old female SPF BALB/c mice and 6 to 8 weeks-old female SPF chickens were purchased from Vital River Laboratory Animal Technology Co., Ltd (Beijing, China). Mice were kept in individually ventilated cages (IVC), and chickens were kept in high efficiency particulate air (HEPA)-filter isolators in the Animal Care Facilities at CAU. Five to six-month-old male Angora ferrets (*Mustela putorius furo*), serologically negative for currently circulating influenza viruses (H1, H3, H5, H7, and H9) and > 1.0 kg (ranging from 1.10 to 1.80 kg) in weight, were purchased from Angora LTD (Jiangsu, China). All ferrets were housed in wire cages placed inside HEPA-filter isolators. All animals were allowed free access to water and a standard chow diet, and provided with a 12-hour light and dark cycle (temperature: 20-25°C, humidity: 40%-70%). All animals used were not involved in any other experimental procedure.

### Generation of viruses by reverse genetics

As described previously [18], recombinant viruses were generated using a plasmid-based reverse genetics system in the genetic background of 17/H9N2, 17/H7N9, and 22/H3N8. Mutations were introduced into the plasmids using PCR-based site-directed mutagenesis. To check for the absence of unintended mutations, all propagated viruses were subjected to complete sequencing.

### Polymerase activity assay

To compare the polymerase activities of different viral ribonucleoprotein (RNP) complexes, a dual-luciferase reporter assay system (Promega, Madison, WI, USA) was employed [25]. The PB2, PB1, PA, and NP gene segments of respective viruses were individually cloned into the pCDNA3.1 expression plasmid. 125 ng of each of the PB2, PB1, PA, and NP plasmids, as well as 10 ng of the pLuci luciferase reporter plasmid and 2.5 ng of the Raniera internal control plasmid, were transfected into 293T cells or DF-1 cells. The cell cultures were then incubated at 33°C or 37°C for 24 hours. The transfected cells were then lysed and analyzed for firefly and renilla luciferase activities using a GloMax 96 microplate luminometer (Promega). The function of ANP32 was examined using a polymerase assay by co-transfection of DKO cells with different ANP32 proteins or empty vector (20 ng) at 33°C or 37°C for 24 hours. All of the experiments were performed independently at least three times.

### Growth kinetics of viruses in A549 and CEF cells

A549 and CEF cells were cultured in 6-well plates and infected in triplicate with indicated viruses at a multiplicity of infection (MOI) of 0.1 (A549) or 0.01 (CEF). After a 2-hour incubation, cells were washed three times with DPBS, and incubated at 37°C in 5% CO_2_. Supernatants were collected at 12, 24, 36, 48, and 60 hours post-infection (hpi), and virus titers were determined by TCID_50_ assays on MDCK cells.

### Quantitative real-time PCR assay

Levels of mRNA and viral RNA (vRNA) were determined in A549 cells and DF-1 cells infected in triplicate with viruses (17/H9N2^627E^, 17/H9N2^627V^, or 17/H9N2^627K^) at an MOI of 0.1(A549 cells) or MOI of 0.001 (DF-1 cells). Total RNA was extracted from infected cells by using TRIzol reagent according to the manufacturer’s instructions (Invitrogen). For the detection of mRNA and vRNA, the oligo(dT) primer and primer uni-12 (5’-AGCAAAAGCAGG -3’) were used respectively to generate cDNAs from 1 μg of total RNA per sample by using Superscript III first-strand synthesis supermix (Invitrogen). The quantitative real-time PCR (qRT-PCR) mixture for each reaction sample consisted of 10 μL of 2× SYBR green PCR master mix (Applied Biosystems), 7 μL of nuclease-free water, 0.5 μL of each primer, and 2 μL of the cDNA template. qPCR was conducted using a 7500 real-time PCR system (Applied Biosystems) with the following program: 1 cycle at 95°C for 10 min, followed by 40 cycles of 95°C for 15 s and 60°C for 1 min. Expression values for each gene, relative to *β-actin*, were calculated by using the 2^-ΔΔCT^ method. Each qPCR sample contained three technical replicates. Primers for the amplification of the *β-actin* and PB2 genes are as follows: forward primer 5’-AGAGCTACGAGCTGCCTGAC-3’ and reverse primer 5’-CGTGGATGCCACAGGACT-3’ for *β-actin*, forward primer 5’-GGAACAGGAATGGACCGACA-3’ and reverse primer 5’-ACTGAGATCTGCATGACCCG-3’ for PB2.

### Mouse challenge studies

Groups of eight mice were anesthetized by intramuscular injection of Zoletil 50 (Zoletil; Virbac SA, Carros, France) at a dose of 20 mg/kg, and inoculated intranasally (i.n.) with 10^6^ TCID_50_ of test virus or PBS in a total volume of 50 μL. At 3 days post-infection (dpi), 3 mice from each group were euthanized, and lung and nasal turbinate tissues were collected for virus titration in MDCK cells by TCID_50_ assay. Lungs of mice infected with the indicated viruses at 3 dpi were collected for histopathology and immunohistochemistry (IHC). The remaining 5 animals were monitored in the remaining two weeks for disease progression, weight change, and mortality. Mice that experienced a weight loss exceeding 25% of their original weight were humanely euthanized.

### Chicken challenge studies

Experimental procedures for virus infection of chickens were under previously described protocol [51]. Groups of 6 chickens were inoculated i.n. with 10^6^ EID_50_ of indicated virus or PBS in a volume of 200 μL. At 3 dpi, 3 chickens from each group were euthanized, and lung and nasal turbinate tissues were collected for virus titration by EID_50_ assay. Lungs of chickens at 3 dpi were collected for histopathology and IHC. The remaining 3 chickens were monitored and tracheal swabs were taken at 2, 4, 6, and 8 dpi for virus titration by EID_50_ assay.

### Histopathology and IHC analysis

Lungs were fixed in 10% buffered formalin, embedded in paraffin, sectioned and stained with hematoxylin and eosin (H&E). Tissue sections were also immunostained with a viral NP monoclonal primary antibody (ab20343, Abcam, Beijing, China). The secondary antibody used was conjugated to horse radish peroxidase (HRP), and the color reaction was based on the use of an HRP reaction kit (ZSGB-BIO, Beijing, China). Histopathological examination was conducted blind by two experienced experimental pathologists.

### Virus passage in chickens

For virus passage in chickens, as depicted in Fig. 4A, groups of 2 SPF chickens were inoculated at passage 1 (P1) with 10^6^ EID_50_ of the indicated virus in a volume of 200 μL. At 3 dpi, 2 naïve chickens were co-housed with the P1 chickens to establish the P2 generation. At 6 dpi of P1 chickens, lung tissues of P1 chickens were collected and homogenized in 500 μL of cold PBS, and another 2 naïve chickens were introduced as P3. The same sequential steps were used for passage up to the P5.

### Ferret transmission experiments

The transmission study consisted of groups of 3 ferrets, following a previously described protocol[32,52] comprising one infected, one direct contact (DC), and one respiratory droplet contact (RD) group. Ferrets were housed in wire cages inside isolators, with each isolator approximately 15 cubic feet in size. Each ferret was sedated with ketamine (20 mg/kg) and xylazine (1 mg/kg) via intramuscular injection and inoculated intranasally with 10^6^ TCID50 of 17/H9N2^627E^, 17/H9N2^627V^, or 17/H9N2^627K^ virus in a volume of 500 μL delivered at 250 μL per nostril. Twenty-four hours later, 2 naïve ferrets were introduced into the isolator, with the direct contact ferret placed in the same cage as the infected ferret, while the respiratory droplet contact ferret was placed in a cage separated by a wire mesh wall to prevent physical contact.

Individual weight was monitored throughout the study. Nasal washes were collected from the ferrets by 2 mL virus transport medium every other day for 14 days and titrated for viruses in MDCK cells by TCID50 assays. Additionally, 200 μL nasal washes were used for RNA extraction and deep sequencing. At 21 dpi, blood samples of ferrets were collected to determine seroconversion using the HI Assay.

### Next generation sequencing

A total of 54 ferret nasal wash samples, 6 chicken lung samples, and 3 viruses (17/H9N2^627E^, 17/H9N2^627V^, and 17/H9N2^627K^) were subjected to whole-genome sequencing. Viral RNA was harvested from the samples by using the QIAamp Viral RNA Mini Kit (Qiagen). Multiplex reverse transcription-PCR amplification of all 8 influenza virus genome segments was performed on RNA samples using PrimeScript™ II 1st Strand cDNA Synthesis Kit (Takara 6210) and primers Uni12/Inf1 (5’-GGGGGGAGCAAAAGCAGG-3’), Uni12/Inf3 (5’-GGGGGAGCGAAAGCAGG-3’), and Uni13/Inf1 (5’CGGGTTATTAGTAGAAACAAGG-3’). As previously described [32], PCR products were purified on spin columns and quantified by using the Quant-It Pico Green kit (Invitrogen, Carlsbad, USA). The purified amplicons were pooled in equimolar and paired-end sequenced (23150) on an Illumina NovaSeq platform according to the standard protocols. Sequencers by 200 or 250 bp paired-end sequencing, and sequencing depth for avian influenza virus isolates was about 0.2 G per sample. NGS findings were validated by qRT-PCR. Raw NGS reads were processed by filtering out low-quality reads (eight bases with quality < 66), adapter-contaminated reads (with > 15 bp matched to the adapter sequence), poly-Ns (with 8 Ns), duplication and host-contaminated reads (SOAP2 [version 2.21]; less than five mismatches). Sanger sequences of parental viruses were chosen as reference sequences. Burrows-Wheeler Alignment Tool (version 0.7.12) and SAMtools (version 1.4) were then used to perform reference-based assembly. Data analyses were performed using a custom quality-based variant detection pipeline.

To detect the genomic variants, the iSNVs were identified as following standards: 1) reads > 1000; 2) mutation rates >1%. The iSNVs frequency was calculated by using the number of iSNVs divided by the total number of genome sites.

### Quantification and statistical analysis

Graphing and statistical analyses were performed using GraphPad Prism 8 (GraphPad Software, San Diego, California, USA; www.graphpad.com). Experimental groups were statistically compared by analysis of variance (ANOVA). P value < 0.05 was considered to indicate statistical significance.

## Acknowledgments

We thank the Global Initiative on Sharing Avian Influenza Data (GISAID) for sharing the influenza virus sequence data used in this study. We are grateful to Xiaojun Wang (Harbin Veterinary Research Institute, China) for providing the DKO cells, Shangang Jia (China Agricultural University, China) for helpful suggestions on data analysis, and Nianzhi Zhang (China Agricultural University, China) for constructive suggestions on structure analysis. We thank BioRender (biorender.com) for assisting with graphics.

## Funding

National Key Research and Development Program of China 2022YFF0802403 (JP)

National Key Research and Development Program of China 2021YFD1800202 (JP)

National Natural Science Foundation of China 32192450 to (JL)

National Natural Science Foundation of China 32102661 (ZJ)

UK Biotechnology and Biological Sciences Research Council BBS/E/D/20002173 (PD and LL)

## Author contributions

Research designing: JP, YG

Data collection and analysis: YG, YZ, YL

Experiment: YG, SS, WP, TL, LW, XC, JD, CZ, FD

Funding acquisition: JP, JL, ZJ, PD, LL

Writing – original draft: YG, JP

Writing – review & editing: JL, HLY, LL, PD, KCC, MHW, HS, YS, ZJ, FD

## Competing interests

The authors declare that they have no competing interests.

## Data and materials availability

Genetic variation data of H9N2 viruses during passage in chickens are available at Zenodo data (DOI: 10.5281/zenodo.12730083). All other data supporting the findings of this study are available in the main text or the supplementary information.

## Notes

### Competing Interest Statement

The authors have declared no competing interest.

### Summary of Updates

We have updated figure S5 in the supplementary files.

