## Supplementary figures and images for "An emerging PB2-627 polymorphism increases the pandemic potential of avian influenza virus by breaking through ANP32 host restriction in mammalian and avian hosts"

### Fig S1

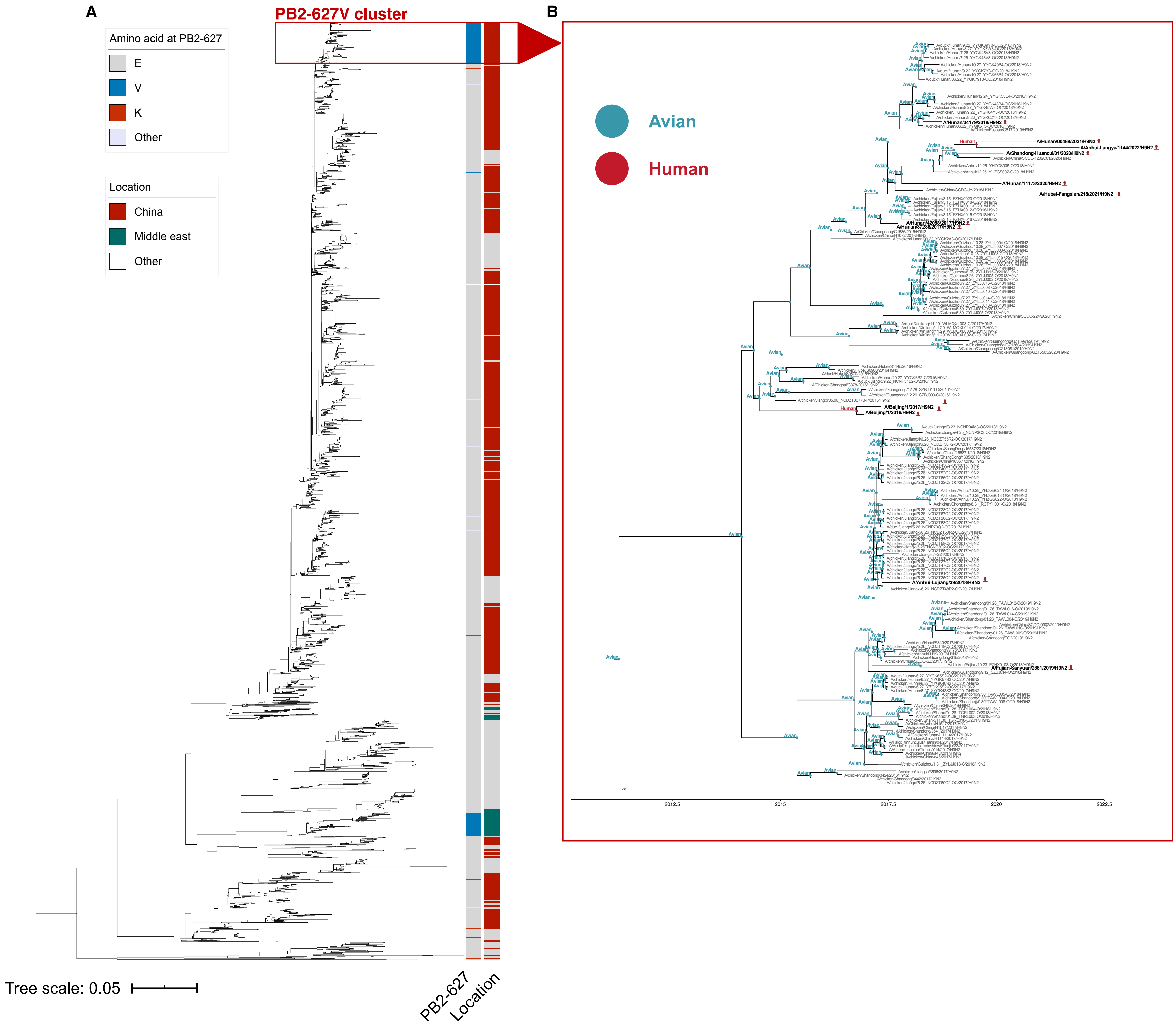
