## Supplement figures and tables for "An emerging PB2-627 polymorphism increases the pandemic potential of avian influenza virus by breaking through ANP32 host restriction in mammalian and avian hosts"

**This PDF file includes:**

S2 to S5 Fig

S1 to S3 Table

**Other supplementary files for this manuscript include the following:**

S1 Fig

Datasets S1

**S1 Fig. H9N2 virus and its reasortants with PB2-627V are in the same cluster. (A).** A total of 4523 H9N2 PB2 sequences were collected from GISAID, Amino acid at PB2-627 residue and location of all isolates are shown. **(B).** The MCC tree of the PB2-627V cluster. The internal nodes indicated the most recent common ancestor of sub-group, which is divided into avian and human. All human strains’ names are in bold.

(separate PDF file)

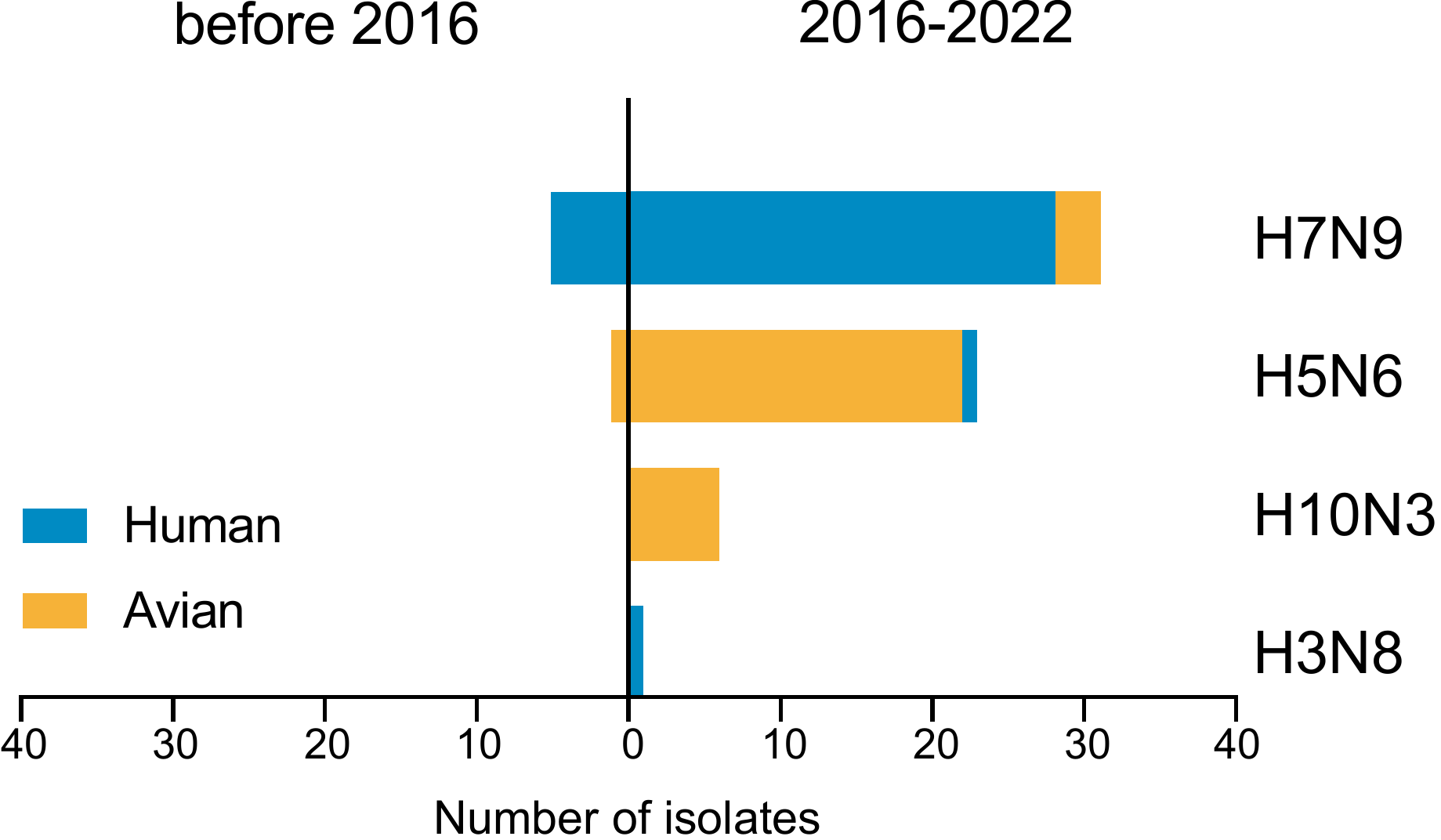

**S2 Fig. Increased detection of PB2-627V in novel AIVs reassortant with H9N2 PB2 gene since 2016.** All the sequences of H7N9, H5N6, H10N3 and H3N8 PB2 genes were obtained from the GISAID database. Viruses with PB2 gene identity greater than 97% to that of PB2 gene of H9N2 viruses were considered reassortants of H9N2. The bars depict the number of isolates with PB2-627V in different subtypes and hosts before and after 2016.

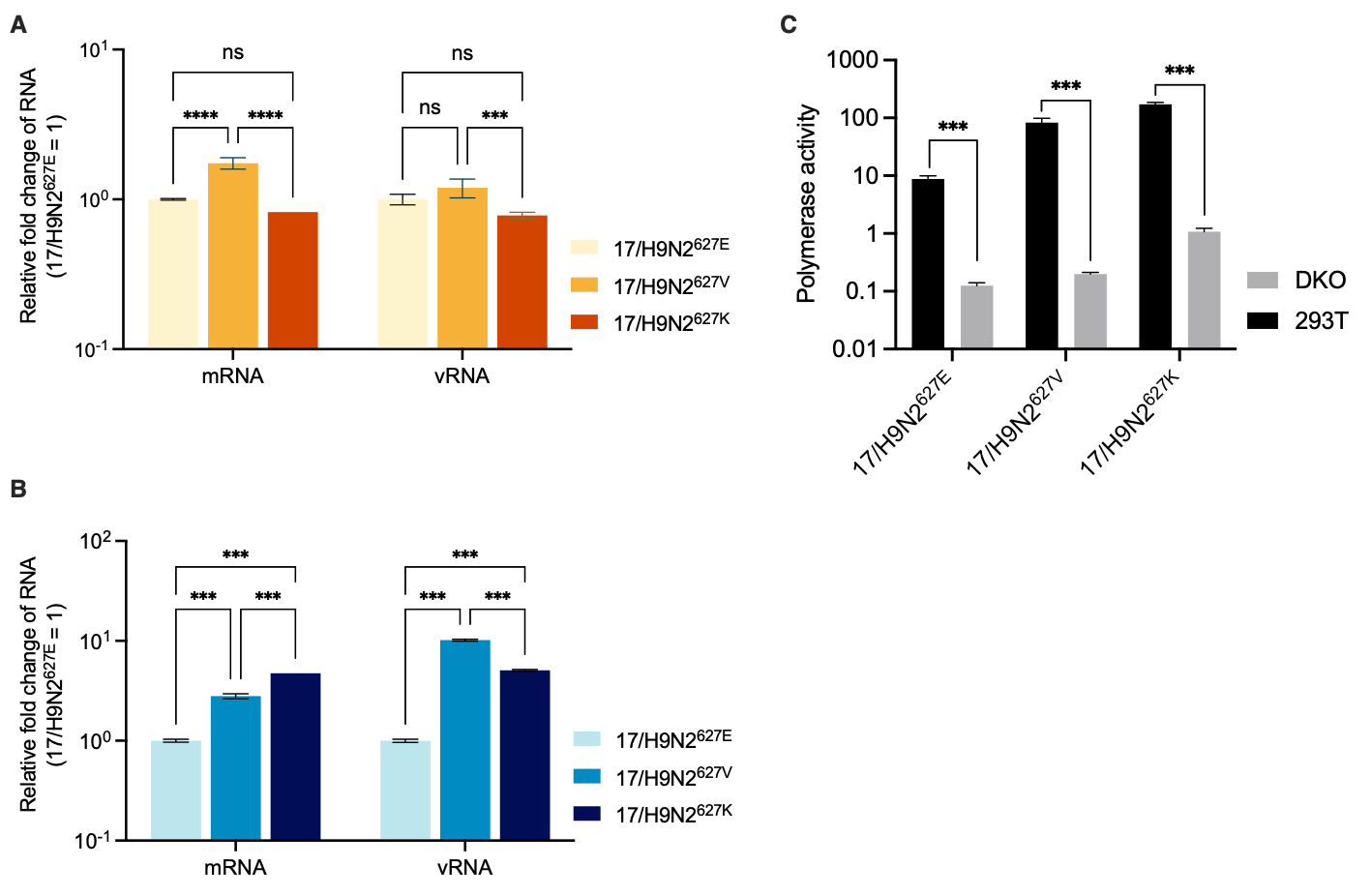
**S3 Fig. Host ANP32 required for viral polymerase and differential viral RNA production in avian and human cells. (A, B).** DF-1 cells **(A)** and A549 cells **(B)** were infected with indicated viruses at MOI of 0.001 and 0.1 respectively. The cells were cultured at 37℃ for 24 hours after infection. PB2 mRNA and vRNA, normalized to β-actin expression, determined by quantitative real-time PCR. **(C).** Wild-type 293T cells and DKO cells were transfected with firefly minigenome reporter, renilla expression control, and polymerase genes of 17/H9N2^627E^, 17/H9N2^627V^, or 17/H9N2^627K^. The cells were cultured at 37℃ for 24 hours after transfection. Data indicate the firefly luciferase gene activity normalized to the renilla luciferase gene activity. Statistical differences between samples are indicated, based on one-way ANOVA with Dunnett’s testing (ns, not significant; *, p <0.05; **, p < 0.01; ***, p <0.001). Error bars are SEM in one representative experiment.

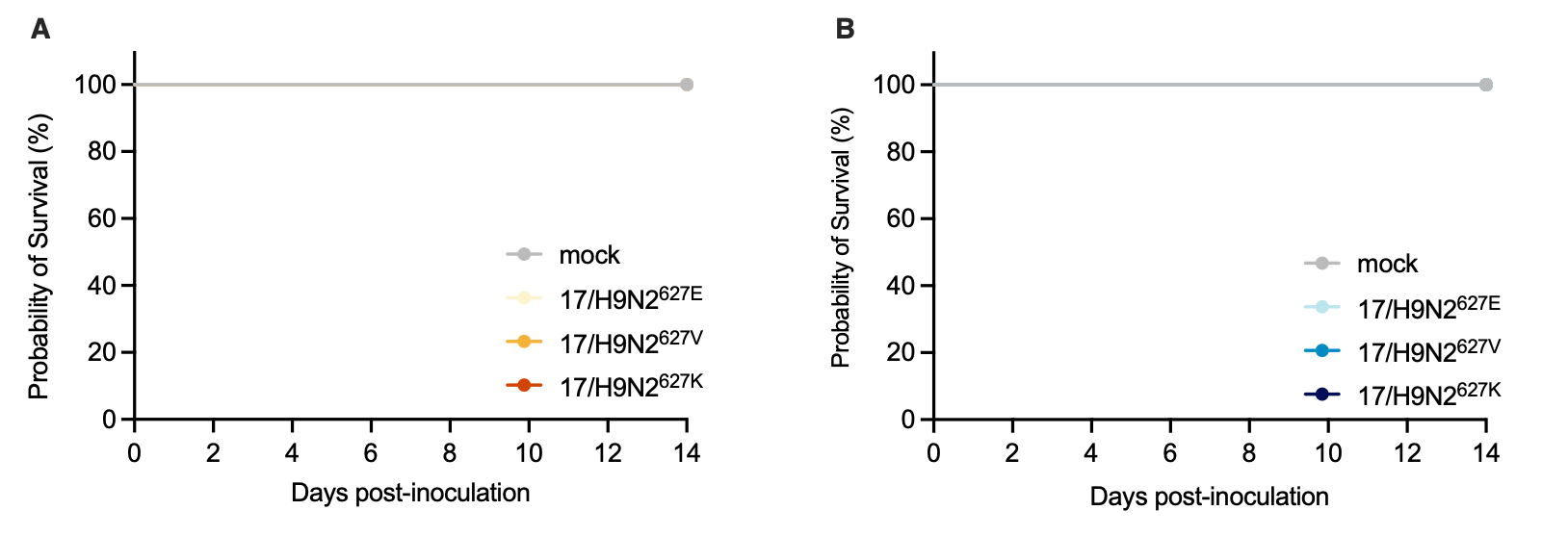

**S4 Fig. Survival of infected mice and chickens.** **(A)**. Groups of 3 chickens were i.n. infected with indicated viruses at 10^6^ EID_50_, and the survival were monitored for 14 days. **(B).** Groups of 5 mice were infected with indicated virus at doses of 10^6^ TCID_50_ and the survival were monitored for 14 days.

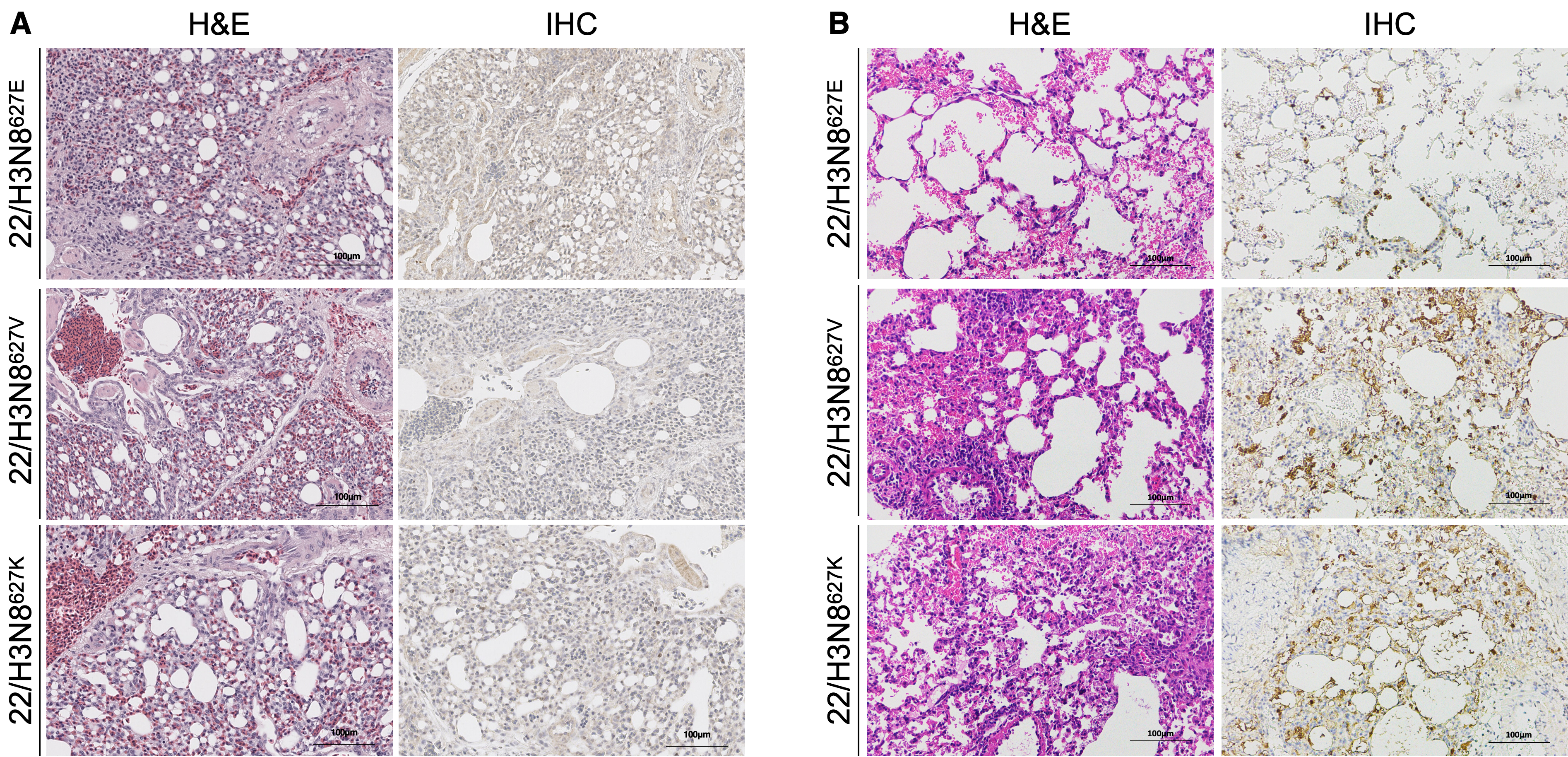

**S5 Fig. H&E and IHC of chicken and mouse lungs infected with PB2-627 variant H3N8 viruses. (A).** Chickens were infected at 10^6^ EID_50_ with indicated viruses, and three chickens per group were euthanized at 3 dpi. **(B).** Mice were infected at 10^6^ TCID_50_ with indicated viruses, and three mice per group were euthanized at 3 dpi. Lung sections were stained with H&E and subjected IHC for the detection of viral NP protein. Scale bars, 100 μm.

**S1 Table. Hosts, geographical distribution, and subtypes of AIVs carrying PB2-627V.**

| Host | Number | Percentage |  | Country | Number | Percentage |  | Subtype | Number | Percentage |
| --- | --- | --- | --- | --- | --- | --- | --- | --- | --- | --- |
| Avian^a^ | 281 | 71.32% |  | China | 243 | 61.68% |  | H9N2 | 254 | 64.47% |
| human | 58 | 14.72% |  | Israel^c^ | 48 | 12.18% |  | H7N9 | 39 | 9.90% |
| swine | 30 | 7.61% |  | Egypt^c^ | 41 | 10.41% |  | H5N6 | 36 | 9.14% |
| Environment | 20 | 5.08% |  | Germany | 18 | 4.57% |  | H1N1 | 18 | 4.57% |
| Red-fox^b^ | 3 | 0.76% |  | Mexico | 16 | 4.06% |  | H5N2 | 18 | 4.57% |
| Tiger | 1 | 0.25% |  | Jordan^c^ | 11 | 2.79% |  | H1N2 | 10 | 2.54% |
| Mink | 1 | 0.25% |  | Canada | 4 | 1.02% |  | H10N3 | 6 | 1.52% |
|  |  |  |  | Italy | 4 | 1.02% |  | H5N1^d^ | 5 | 1.27% |
|  |  |  |  | Spain | 3 | 0.76% |  | H3N2 | 3 | 0.80% |
|  |  |  |  | Russia | 3 | 0.76% |  | H3N8 | 3 | 0.76% |
|  |  |  |  | America | 1 | 0.25% |  | H6N6 | 1 | 0.25% |
|  |  |  |  | Denmark | 1 | 0.25% |  | H7N1 | 1 | 0.25% |
|  |  |  |  | Belgium | 1 | 0.25% |  |  |  |  |
| ^a^. Avian species include chicken, duck, turkey, goose, quail, mallard, and wild bird. ^b^. All 3 red fox isolates are 2.3.4.4b H5N1 viruses. ^c^. These countries are in the Middle East.  ^d^. One H5N1 virus belongs to 2.3.4.4e clade; four H5N1 viruses belong to 2.3.4.4b clade. | | | | | | | | | | |

**S2 Table. Clinical symptoms of ferrets infected with H9N2 influenza viruses**

| Virus | Maximum body weight loss (%) | | |  | Seroconversion (HI^a^ antibody titer range) | | | Respiratory droplet transmission |
| --- | --- | --- | --- | --- | --- | --- | --- | --- |
|  | Donor | DC | RD |  | Donor | DC | RD |  |
| 17/H9N2^627E^ | 5.89 | 8.00 | 2.00 |  | 3/3  (512-1280) | 3/3  (512-1280) | 0/3 | None |
| 17/H9N2^627V^ | 9.94 | 6.19 | 5.98 |  | 3/3  (256-1280) | 3/3  (512-2560) | 3/3  (1280-1280) | 3/3 |
| 17/H9N2^627K^ | 8.00 | 4.35 | 6.80 |  | 3/3  (256-1280) | 3/3  (512-2560) | 3/3  (512-2560) | 3/3 |
| ^a^. HI, hemagglutination inhibition | | | | | | | | |

**S3 Table. Area under the curve (AUC) values of virus output of infected/exposed ferrets**

| Virus | Donor | | |  | DC | | |  | RD | | |
| --- | --- | --- | --- | --- | --- | --- | --- | --- | --- | --- | --- |
|  | 1 | 2 | 3 |  | 1 | 2 | 3 |  | 1 | 2 | 3 |
| 17/H9N2^627E^ | 16.38 | 11.50 | 11.38 |  | 17.63 | 13.00 | 5.98 |  | 0.00 | 0.00 | 0.00 |
| 17/H9N2^627V^ | 18.38 | 16.75 | 11.88 |  | 20.00 | 20.50 | 13.88 |  | 9.50 | 10.75 | 15.00 |
| 17/H9N2^627K^ | 16.38 | 10.25 | 11.63 |  | 16.50 | 17.38 | 14.88 |  | 17.25 | 14.75 | 14.25 |
